# An off-the-shelf multi-well scaffold-supported platform for tumour organoid-based tissues

**DOI:** 10.1101/2022.08.14.503643

**Authors:** Nancy T Li, Nila C Wu, Ruonan Cao, Jose L Cadavid, Simon Latour, Xiaoya Lu, Yutong Zhu, Mirjana Mijalkovic, Reza Roozitalab, Alison P McGuigan

## Abstract

Complex 3D bioengineered tumour models provide the opportunity to better capture the heterogeneity of patient tumours. Patient-derived organoids are emerging as a useful tool to study tumour heterogeneity and variation in patient responses. Organoid cultures typically require a 3D microenvironment that can be manufactured easily to facilitate screening. Here we set out to create a high-throughput, “off-the-shelf” platform which permits the generation of organoid-containing microtissues for standard phenotypic bioassays and image-based readings. To achieve this, we developed the Scaffold-supported Platform for Organoid-based Tissues **(SPOT)** platform. SPOT is a 3D gel-embedded *in vitro* platform that can be produced in a 96- or 384-well plate format and enables the generation of flat, thin and dimensionally-defined microgels. SPOT has high potential for adoption due to its reproducible manufacturing methodology, compatibility with existing instrumentation, and reduced within-sample and between-sample variation, which can pose challenges to both data analysis and interpretation. Using SPOT we generate cultures from patient derived pancreatic ductal adenocarcinoma organoids and assess the cellular response to standard-of-care chemotherapeutic compounds, demonstrating our platform’s usability for drug screening. We envision 96/384-SPOT will provide a useful tool to assess drug sensitivity of patient-derived organoids and easily integrate into the drug discovery pipeline.

## 1.0 Introduction

Cancer is a family of complex diseases that result from its heterogeneous and dynamic nature [1]–[3]. Although substantial progress has been made toward understanding the disease, cancer still accounts for one in seven deaths worldwide [4]. This can partly be attributed to the high failure rate of new anti-cancer drug candidates due to the lack of predictive models used in the pre-clinical phases of the drug discovery pipeline [5]–[7]. Historically, *in vitro* phenotypic studies for drug screening are commonly performed on two-dimensional (2D) monolayers where a homogeneous population of tumour cells (i.e. an immortalized cell line) is exposed to potential drug candidates. In combination with typically soluble assays of cell viability, this method enables rapid data acquisition and interpretation, leading to widespread adoption by the cancer research community. However, these models lack the complex biophysical and biochemical cues presented by the tumour microenvironment (TME). To address the need for *in vitro* models which better mimic features of the TME, such as its three-dimensional (3D) architecture and naturally derived extracellular matrix (ECM), many engineered *in vitro* models for screening anti-cancer drug candidates have been proposed, such as microfluidic organ-on-a-chip models [8], [9], organotypic models [10], [11], or multicellular spheroids [12]–[14]. However, there has generally been low adoption of complex 3D bioengineered models beyond the bioengineering research community, due to technical challenges associated with the experimental setup, including lack of assay flexibility, robust readouts, reproducible manufacturing techniques, and scalability [15]. As a result, only simplistic easy-to-adopt models are common, such as 3D spheroid models [16]–[18] and gel-embedded microtissues such as hydrogel domes or plugs [16], [19], [20]. A number of tools have emerged to generate arrays of spheroids or microtissues either in suspension [16] or encapsulated in hydrogel ECM materials [14], [16], [21] with some commercial success [22].

More recently, there has been a paradigm shift toward the use of cancer patient-derived organoid (PDO) models which recapitulate morphological characteristics of the primary tissue [8], [23]–[25] and better capture the patient heterogeneity that has been shown to impact cancer progression, prognosis, and drug sensitivity [26]–[28]. The incorporation of these specialized patient-derived cell types within *in vitro* models has the potential to allow phenotypic studies of complex biological processes in heterogeneous cell populations, to improve the translation of drug candidates to the clinic. Approaches to screen the impact of various drugs on different organoid models to explore patient heterogeneity have primarily cultured organoids in 2D monolayers [29], [30], on the surface of hydrogel layers with dilute hydrogel in the surrounding media (the “slurry” method) [17], or within 3D hydrogel plugs [31]. However, only the hydrogel plug model creates the 3D culture conditions that are typically required by PDOs (i.e., to enable self-organization) [32]. Therefore, suspension of the PDOs in hydrogel plugs has emerged as the highest throughput approach to enable 3D drug screening [24], [31], [33].

A technical drawback of the hydrogel plug model however, is the curved gel surface that forms in a standard well plate due to meniscus effects [25], requiring higher sample volumes to ensure complete coverage of the well bottom, thus reducing the ratio of media volume to tissue ratio and requiring more frequent cell medium replenishments to maintain the microtissue **(SI Table 1)**. The uneven shape of the gel surface also presents challenges for image-based acquisition and analysis, consequently limiting assay readouts to soluble measures of cell viability. Alternatively, a hydrogel dome can be used to reduce the volume of gel required, however dome culture cannot be easily created in a small well size (for example, a 384-well plate) without collapsing, cannot be easily automated for assay scaling, and still presents imaging challenges as the gel surface is also curved. A need exists therefore for a 3D hydrogel culture platform compatible with PDO cultures, with the potential for high-throughput manufacturing and that facilitates high-throughput phenotypic (likely image-based) analysis which has been shown to acquire richer information about cellular behaviour[34]. In previous work, we reported a 3D *in vitro* model, Gels for Live Analysis of Compartmentalized Environments (GLAnCE), in which cells are suspended in a thin layer of gel to enable non-destructive data acquisition over time using widefield microscopy, rather than time-laborious and data-intensive confocal microscopy [35], [36]. While GLAnCE addresses the need for a 3D culture platform that enables ease-of-imaging, the throughput possible with GLAnCE is limited because it requires multiple assembly steps (i.e., pre- and post-seeding) and would be challenging to integrate with automated robotic manufacturing.

An alternative strategy to create a meniscus-free gel-embedded organoid culture platform that is more amenable to high-throughput construction, is the use of a scaffold component to reinforce and mold the hydrogel structure. Paper scaffolds have been used to create paper-stacked models by us [37]–[42] and others previously [43]–[46], to reinforce hydrogel structures in which the fibers form a complex porous network which act as anchor points for cellular adhesion and provides mechanical strength for easy sample handling. These paper-based models have been used to probe invasive behaviour [41], [42], [47]–[50], metabolomic gradients [38], [51], drug-induced cytotoxicity [44], [45] and radiation-induced cytotoxicity [52]. Typically, these paper-based culture models are fabricated by cutting individual disks of Whatman® filter paper and placing these disks within a well-plate [50], [53], [54]. Throughput can be improved by adapting techniques designed for analytical devices [55]–[58], such as paper-based ELISA [59], to generate a continuous scaffold array with individual wells which are micropatterned using photolithography techniques [60]–[62], thermal lamination of polymer films [48], [52], [63] or wax-printing [44], [45], [64], [65]. However, many of these previously reported paper-based platforms have relied on fluorescent gel scanners to measure cytotoxic activity at a defined assay endpoint, which required the device to be disassembled and the paper scaffold to be removed from the well plate for analysis. The complexity of these designs leads to a requirement for manual assembly and disassembly, while also limiting the ability to capture time-course cellular dynamics *in situ*. This ultimately limits the overall experimental throughput achievable using these designs.

Here we set out to create a well-plate paper-based device for organoid culture that could be fabricated for “off-the-shelf” use, with potential for high-throughput studies using data acquisition pipelines which avoid destruction of the microtissues for analysis, which is important for studies where cellular dynamics are monitored over time. In this report we describe the development of a paper-embedded multi-well system named SPOT (**S**caffold-supported **P**latform for **O**rganoid-based **T**issues) which can be produced in both a 96-well (96-SPOT) and a 384-well (384-SPOT) format. By exploiting the high permeability of the cellulose scaffold, we infiltrate low volumes of gel-cell material into a prefabricated well-plate to generate tumour microtissues, therefore providing an “off-the-shelf” device. Further, 96-SPOT and 384-SPOT have high potential for adoption due to their reproducible manufacturing and seeding methodology, and capability to monitor cellular dynamics in a format compatible with standard instrumentation. We also integrate PDOs into the platform and demonstrate we can test the drug sensitivity of PDOs exposed to five monotherapies commonly used in the treatment of pancreatic ductal adenocarcinoma (PDAC). We envision that 96/384-SPOT can address the current need to assess the drug sensitivity and growth dynamics of PDOs in order to accelerate the drug discovery pipeline.

## 2.0 Materials and Methods

### Cell culture

The KP4 pancreatic tumour cell line (JCRB Cellbank) was maintained in IMDM containing 10% fetal bovine serum (FBS) (Fisher Scientific, New Hampshire, USA), and 1 μg/ml penicillin and streptomycin (Sigma-Aldrich, Canada). Medium was changed every two to three days and the cells were passaged every three to four days. Pancreatic ductal adenocarcinoma organoids established from PDAC patients were obtained from the UHN Biobank at Princess Margaret Cancer Centre (PMLB identifier PPTO.46 from the University Health Network, Ontario, Canada) under a protocol in compliance with the University of Toronto Research Ethics Board guidelines (protocol #36107). Organoid cultures were maintained in Advanced DMEM/F-12 (Gibco, ThermoFisher Scientific, Waltham, USA) supplemented with 2 mM GlutaMAX, 10 mM HEPES (Gibco), 1% penicillin/streptomycin, 1X W21 supplement (Wisent), 1.25 mM N-Acetyl-L-cysteine, 10 nM Gastrin I (1-14) (Sigma Aldrich, St. Louis, USA), 50 ng/mL recombinant human EGF, 100 ng/mL recombinant human noggin, 100 ng/mL recombinant human FGF-10 (Peprotech, Rocky Hill, USA), 0.5 μM A83-01 (Tocris Biosciences, Bristol, United Kingdom), 10 μM Y-27632 (Selleck Chemicals, Houston, USA), 10 mM nicotinamide (Sigma Aldrich), 20% v/v Wnt-3a conditioned media and 30% v/v human R-spondin1 conditioned media (Princess Margaret Living Biobank, Ontario, Canada). PPTO.46 cells were cultured in 48-well polystyrene plates in 40 μL domes of Growth Factor Reduced Phenol Red-Free Matrigel™ Matrix (Corning Life Sciences, Corning, USA) with 500 μL of complete media. Media was refreshed twice per week and PPTO.46 cells were passaged once per week (1:8 split ratio). PDOs were used for a maximum of 40 passages. All cultures were maintained in a humidified atmosphere at 37 °C and 5% CO_2._

### Lentivirus production and cell transduction

Cells were tranduced to express green fluorescence protein (GFP). Lentivirus was produced using calcium phosphate co-transfection of HEK293T cells with the psPAx2 plasmid (packaging vector, Addgene #12260), pMd2g plasmid (VSVG envelope, Addgene #12259) and the pLenti-CMV-GFP-Puro plasmid (Addgene #17448) as previously described [66]. Supernatants containing viral particles were harvested 48 h after transfection and concentrated using a 20 mL 100,000 MWCO spin column (Vivaspin ®, Cytiva, USA). Wild-type KP4 cells were transduced with concentrated virus and GFP-expressing cells were selected using puromycin selection (1 μg/mL, Sigma-Aldrich, St Louis, USA). GFP-expressing PDOs were created using the same virus as KP4 but with a slightly different transduction protocol. PDOs were suspended in infection media and spinoculated for 1 hour at 600 g at 32 °C, before being resuspended and incubated for an additional 6 hours at 37 °C in 5% CO_2_. PDOs were then washed and seeded into Matrigel™ with fresh media. 48 hours after infection, cells were sorted for the middle 80% of the GFP+ population.

### Multi-well plate component fabrication and assembly

Cellulose scaffolds (Miniminit Products, R10, Scarborough, Canada) were partially infiltrated with a poly(methyl methacrylate) (PMMA, 120KDa, Sigma-Aldrich, St. Louis, USA) solution in acetone (Sigma-Aldrich, St. Louis, USA) to delineate the inter-well regions from the area available for cell infiltration. The PMMA-acetone solution (1.75g PMMA to 7.84mL acetone) was mixed with blue nail polish (Essie) for visibility. A modification of the contact wicking printing technique developed in our group [39], [40] was used for PMMA infiltration. Briefly, the scaffold templates were taped to a sheet of parchment paper to prevent sticking and placed on the AxiDraw V3 (Evil Mad Scientists, USA) plotter support. The AxiDraw V3, a commercial 2D plotter, is controlled by Inkscape (https://inkscape.org/) with an AxiDraw extension (https://wiki.evilmadscientist.com/Axidraw_Software_Installation), which was used to move the printhead, which contained a 5 mL syringe loaded with 0.5 mL of PMMA-acetone solution, in the desired array of 96 rings with a nominal diameter of 7 mm to match the 96-well no-bottom well plate (Greiner, catalog #82050-714). To further cover the surrounding inter-well regions, an opensource traveling salesperson pattern (TSP) generator was used (https://mitxela.com/plotterfun/). An Inkscape pattern of the 96-well array was saved as a PNG image and imported to the mixtela generator. Using the ‘stipples’ algorithm, a TSP pattern was generated with maximum brightness set to 200, and maximum stipples set to 5000. After rendering, a SVG file is exported and layered onto the pattern of the 96-well array. The solution was dispensed through a 20-gauge standard Luer Lock tip (McMaster-Carr, Elmhurst, USA) in constant contact with the cellulose scaffold. As the printhead moved, PMMA slowly wicked into the paper creating a line in the direction of movement; acetone quickly evaporated leaving behind a sheet of PMMA that blocks the pores in the scaffold and makes it impermeable. For the 384-well pattern, the AxiDraw V3 plotter path was controlled directly using an interactive python API (https://axidraw.com/doc/py_api/#introduction) to create a grid pattern corresponding to a 384-well no-bottom well plate (Greiner, catalog # 82051-262). For the 384-well array, the solution was dispensed through a 16-gauge standard Luer Lock tip (McMaster-Carr, Elmhurst, USA). For each plate, two sheets of double-sided, poly-acrylic adhesive tape (Adhesive Research, ARcare, catalog # 90106NB) were cut with an array of 96 circular holes or an array of 384 square holes corresponding to the no-bottom 96 well plate or no-bottom 384 well plate, respectively, using the Silhouette Cameo Electronic Cutting Machine (Silhouette, USA). All components were UV sterilized and assembled in a sterile environment. The double-sided tape was applied to the no-bottom well plate, followed by the PMMA-patterned scaffold sheet, and another layer of double-sided tape. Lastly, a thin, transparent polycarbonate film (McMaster-Carr, catalog # 85585K102) was cut to 110 mm x 75 mm and attached to the bottom of the well plate. An alignment guide was created for the 384-well pattern to guide the device assembly. The assembled well plates were clamped prior to use to prevent delamination of the layers.

### Assessment of device manufacturing robustness

To assess variation between PMMA printed scaffold sheets, printed patterns were scanned with a Cannon MX320 document scanner as 600 dpi tiff images. Images were imported into the open-source image analysis software Fiji (https://imagej.net/Fiji/Downloads) and then we extracted radial (i.e. 96-well pattern) or linear (i.e. 384 well pattern) dimension measurements by thresholding the binary images of the printed patterns. This data was used to assess print pattern variation. To ensure complete polymer infiltration of the cellulose scaffold we performed scanning electron microscopy (SEM). SEM images were obtained using a Hitachi SEM SU3500 (Hitachi High-Technologies Canada Inc., Toronto, Canada). Samples were mounted onto carbon-tape coated stubs and gold–palladium sputter coated for 55 s using a Bal-Tec SCD050 Sample Sputter Coater (Leica Biosystems, USA) and then imaged at 5 kVD. To assess leakage between wells in the assembled multwell plate device the movement of a fluorescence dye between adjacent wells was assessed. Fluorescein solution (Sigma-Aldrich, St. Louis, USA) was added to alternating wells in a checkerboard pattern, with PBS in the remaining wells. The fluorescence intensity was measured using a microplate reader (Tecan Infinite Pro 200 MPlex) at 0 h and 24 h.

### Cellulose scaffold seeding

Collagen hydrogel was prepared by mixing eight parts type I bovine collagen (PureCol 3 mg ml−1; Advanced BioMatrix) with 1 part 10x minimal essential medium (MEM, Life Technologies, Grand Island, USA) by volume and neutralizing to the pH 7 endpoint with 0.8M NaHCO3 (Sigma-Aldrich). The solution was kept on ice prior to addition of cells. Following a standard trypsinization protocol, the adherent cells (i.e. KP4 cells) were pelleted by centrifugation (300*g*, 5 min, 4 °C) and resuspended in an appropriate volume of collagen to achieve the desired cell concentration. In the case of organoids, the PPTO.46 cells were collected by mechanical disruption of the Matrigel™ domes followed by a 10 min incubation in TrypLE Express (Gibco) at 37 °C. This process was repeated until mostly single cells were observed under the microscope (a total of 3 times). PPTO.46 cells were then passed through a 40 μm cell strainer to remove large cell clusters prior to seeding. The remaining cells were pelleted by centrifugation (300*g*, 5 min, 4 °C). In accordance with [39] the PPTO.46 cell pellet was re-suspended in an appropriate volume of a hydrogel blend to achieve the desired cell concentration. The hydrogel blend used for PPTO.46 cells was comprised of 25% Matrigel™ and 75% collagen hydrogel as reported previously [39].

For 96-SPOT experiments, a single-channel micropipette (Gilson) was used to deposit cell-gel from an Eppendorf to the well plate of interest. Prior to use, 70 µL of sterile PBS was added to each of the inter-well spaces in the 96-well plate to prevent evaporation while seeding, and the sterile well plate was placed in a closed and insulated icebox with 4 ice packs for 30 minutes to ensure that all surfaces were cold at the time of seeding. 5 µL of cell-gel was added to the center of each well by carefully dispensing the gel above the paper surface, then gently touching the tip of the pipette to the paper surface to facilitate homogeneous infiltration of the cell-gel in the paper scaffold. For the 384-SPOT, there are no inter-well spaces for the addition of sterile PBS. To minimize evaporation of the gel during seeding, the well plate was seeded within a closed container containing a pre-wet, sterile paper-towel (i.e., a humidification chamber) using an 8-channel electronic micropipette (Gilson, model no. P8X10M-BC) and a plate membrane (Sigma-Aldrich, cat no. Z380059) was used to cover each column of wells as they are seeded. Cell-gel was first dispensed into an 8-well PCR strip. The electronic micropipette was used to dispense 1.5 µL of cell-gel in a column of wells by moving to the center of each well, dispensing the gel above the paper surface, then gently touching the tip of the pipette to the surface of the paper scaffold. For both 96/384-SPOT, after seeding the final well, the well plate was allowed to incubate for 1-2 minutes on the cold ice pack before it was placed in a 37 °C incubator for 45 minutes for hydrogel polymerization. Post-gelation, 200 µL or 60 µL of media was added to each 96-well or 384-well SPOT, respectively.

### Analysis of cell seeding reproducibility

GFP-expressing cells fluorescence was used to assess the variation in seeding between and within wells. Widefield images of GFP-fluorescence in each well were acquired at 4X with an ImageXpress Micro (Molecular Devices, USA) high-content imaging system using both laser-based and image-based autofocus settings (16-bit). Sixteen or four sites were acquired per 96- or 384-SPOT, respectively, to capture a full well bottom, which was subsequently stitched using MetaXpress software (Molecular Devices, USA). A custom script (adapted from [39]) was written in ImageJ to measure the mean grey value (MGV) of GFP signal of 100 randomly selected squares to obtain the standard deviation within 1 well. Standard deviations were used to calculate the coefficient of variation associated to each well using the following:

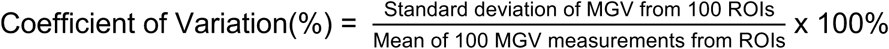

### Measurement of cell metabolism using the alamarBlue® assay

The metabolic quantification alamarBlue® assay (Sigma), following the manufacturer’s instructions, was used to assess cell number variation between culture wells for different cell seeding densities. Briefly, a calibration curve was created for KP4 cells in 96-SPOT over a range of cell densities (0 - 6.0 × 10^7^ cells/mL collagen I hydrogel). This determined an optimal timepoint of 3h for subsequent experiments (data not shown). Fluorescence intensity was measured at a 3h timepoint in a microplate reader at 560-590 nm. Subsequent alamarBlue® experiments to assess variation in cell number between wells were performed in the 96-well SPOT using 10% alamarBlue® reagent in 200 µL per well, or 60 µL per 384-SPOT, with negative control wells containing no cells and positive control wells containing fully-reduced 10% alamarBlue reagent®.

### Quantification of PPTO.46 cell growth in 96-SPOT

GFP-expressing PPTO.46 cells were seeded as single dissociated cells at a cell density of 3 × 10^6^ cells/mL of 25% Matrigel™ and 75% collagen hydrogel. Images of the cultures were taken using the ImageXpress Micro (Molecular Devices, USA) on days 0, 2, 4, and 6 to visually verify that cell growth was linear over 6-days of culture. The MGV of GFP signal was measured for each microtissue using Image J, showing that the cell growth is linear over 6-days of culture.

### Assessing response to PDAC chemotherapeutics

GFP-expressing KP4 cells were plated in 96-SPOT at a cell density of 3 × 10^6^ cells/mL, 10 × 10^6^ cells/mL, and 30 × 10^6^ cells/mL and treated with a 10-fold gemcitabine serial dilution from 0.001 µM to 100 µM to identify an appropriate concentration range which captures 0-100% viability using the alamarBlue® assay. The 10 × 10^6^ cells/mL, and 30 × 10^6^ cells/mL conditions were overpopulated in the low gemcitabine concentration or untreated conditions. All subsequent experiments in SPOT were therefore performed using the cell density of 3 × 10^6^ cells/mL collagen-Matrigel hydrogel.

GFP-expressing KP4 cells were plated at a cell density of 3 × 10^6^ cells/mL in collagen hydrogel (i.e., 15 × 10^3^ cells per 96-SPOT well). Each well was imaged and analyzed to verify homogeneous cell seeding after 1 hour and after 24 hours of incubation. At 24 hours after seeding, 200 µL of media containing Gemcitabine hydrochloride (Sigma-Aldrich, St. Louis, USA) was added to each well, with a 3.16-fold (or half-log) serial dilution that ranged from 3.16 ×10^−3^ µM to 100 µM. Gemcitabine was dissolved in DMSO and was tested in triplicate with a 10% DMSO vehicle control. After 3 days of Gemcitabine treatment, each well was imaged using the ImageXpress Micro (Molecular Devices), then cell viability was also assessed using the alamarBlue® assay (see above).

GFP-expressing PPTO.46 cells were plated at a cell density of 3 × 10^6^ cells/mL in 25% Matrigel™ / 75% collagen hydrogel (i.e., 15 × 10^3^ cells per 96-well or 4.5 × 10^3^ cells per 384-well). Each well was imaged and analyzed to verify homogeneous cell seeding after 1 hour and after 24 hours of incubation. At 24 hours after seeding, 200 µL of media containing the therapeutic compound was added to each 96-well, or 60 µL for each 384-well. Therapeutic compounds were added 24 hours after seeding, after verifying the reformation of organoids. Chemotherapeutics were tested in triplicate using 3.16-fold serial dilutions of gemcitabine and oxaliplatin ranged from 1.0 × 10^−3^ µM to 31.67 µM. Paclitaxel, 5-fluorouracil, and irinotecan (SN-38) ranged from 3.16 × 10^−3^ µM to 100 µM. All compounds were dissolved in DMSO. Vehicle controls were generated corresponding to the DMSO content in the high concentration condition of each respective chemotherapeutic agent. 1 µM staurosporine (Abcam) was used as a 0% viability control. After 5 days of treatment with the respective drug, each well was imaged using the ImageXpress Micro (Molecular Devices), then cell viability was assessed using the alamarBlue® assay.

### Statistics

Statistical analysis was carried out in GraphPad Prism 9 (GraphPad Software, San Diego, California USA). Ordinary one-way ANOVA was used for assessing reproducibility of microtissues and Student t-test was used for comparing IC_50_ values of treated KP4 microtissues or PPTO.46 microtissues. p < 0.05 was considered significant. Superplots were created as described in [67].

## 3.0 Results

### 3.1 Development of a multi-well paper-supported culture platform manufacturing strategy

We set out to develop a multi-well culture platform, Scaffold-supported Platform for Organoid-based Tissues (SPOT), to enable creation of 3D microtissues for imaging of cellular dynamics in a PDAC model. Further, the SPOT platform must utilize inexpensive components, minimize user-dependent assembly processes, be compatible with standard instrumentation, and allow non-destructive *in situ* image-based analysis. The basic design concept of SPOT was to fabricate a multi-well single paper-layer well plate in which the paper layer is sandwiched between two layers of biocompatible double-sided tape to integrate with a device bottom, consisting of a thin (0.127 mm) polycarbonate sheet which was selected to be compatible with microscopy, and a commercial no-bottom well plate to create individually addressable wells **(Figure 1A)**. We selected a commercially available no-bottom well plate to ensure that the resulting well plate would be compatible with a commercial microplate reader using standard settings, or an automated microscope without the need for any major customization. Further, we selected an optically transparent material for the well bottom, to ensure the device would enable image-based readouts of cellular behaviour over time.

**Figure 1:**
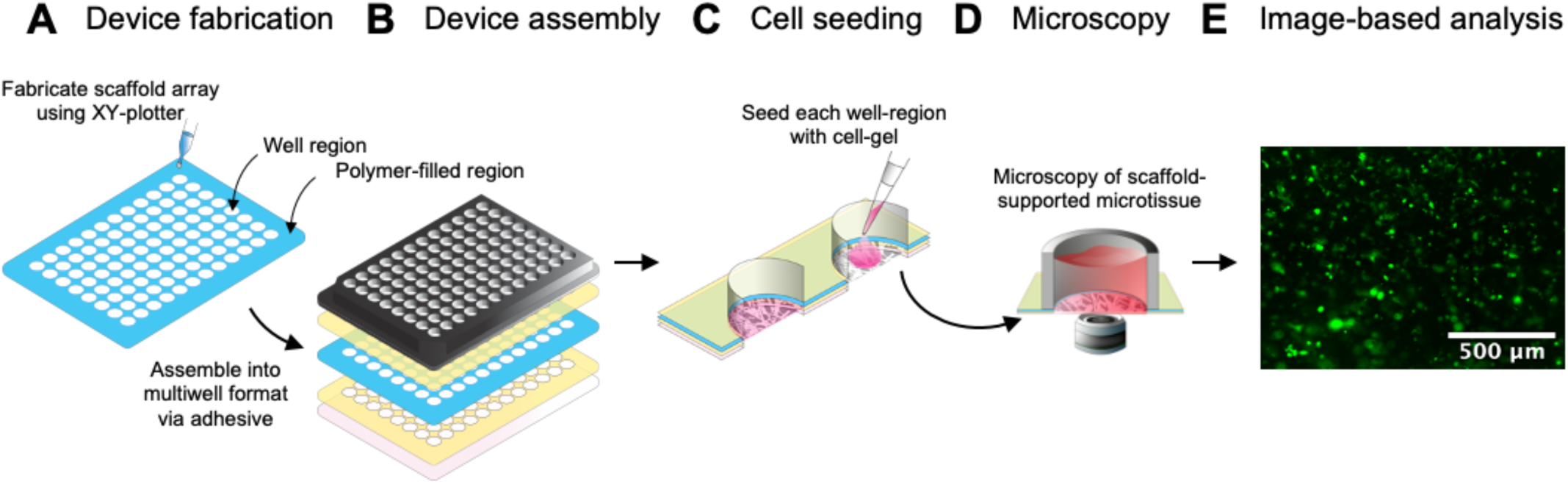
Overview of 96-SPOT device and image analysis workflow. Schematic representation of the experimental workflow. **A)** Printing of well-plate pattern: An XY-plotter is used to infiltrate polymer into a scaffold sheet to generate exposed well regions (white) and polymer-filled regions (blue) **B)** Device assembly: the device is assembled by collating a commercial bottomless well plate (top), PMMA-patterned scaffold (shown in blue), and polycarbonate film (bottom) using layers of double-sided polyacrylic adhesive (shown in yellow) **C)** Cell seeding into scaffold well: cells are suspended in a non-polymerized hydrogel (shown in pink) and pipetted into each well where it infiltrates the scaffold. The hydrogel is then allowed to gel to encapsulate the cells within the scaffold-supported hydrogel **D)** Microscopy: the resulting microtissue can be imaged using automated widefield microscopy, and **E)** Image-based analysis: Acquired images of fluorescent cells (green) suspended in gel can be analyzed using standard image-analysis pipelines. Representative epifluorescence microscopy image of GFP-expressing KP4 cells in one 96-SPOT well. Scalebar is 500 µm.

A cell-gel suspension can then be infiltrated into the paper sheet corresponding to the well regions and gelled to form thin scaffold-supported microgels **(Figure 1B)** which can be imaged **(Figure 1C)** and analyzed easily **(Figure 1D)**. The porous scaffold sheet enabled us to generate dense, gel-impregnated tissues which were mechanically robust due to reinforcement by the scaffold fibers as we have reported previously [37]. The paper scaffold’s thickness also defined the exact dimensions of the microgel enabling the production of thin and flat microgel structures. Specifically, the scaffold provided anchor points for cellular and hydrogel adhesion, thereby minimizing hydrogel contraction issues associated with tissues constructed from naturally derived hydrogels and allowing the generation of dense tissue constructs which more closely resemble the density of *in vivo* tissue [37], while maintaining specific dimensions. The use of the scaffold also significantly reduced the volume of cell-gel and/or reagents required to manufacture the microtissue, when compared to the commonly used hydrogel plug **(SI Table 1)**.

To ensure no liquid transfer between wells, we developed a strategy to pattern a single scaffold sheet to contain an array of isolated wells. Previously, this has been achieved using wax printing [44], [45], [64] and thermal lamination [48], [52], [63]. However, wax exhibited poor adhesion to the double-sided tape, which resulted in a high device failure rate upon plate assembly. We also attempted thermal lamination of polymer films such as polystyrene but found this process was not reproducible. We therefore adapted a contact-capillary wicking printing technique we described previously to print poly-methyl methacrylate (PMMA) into regions of the cellulose scaffold [39] to delineate individual wells in the paper sheet. Specifically, we drew an array with 96 circles **(Figure 2Ai)**, or a grid of lines to create 384 squares **(Figure 2Aii)** using a basic commercially available XY-plotter to move a standard syringe loaded with PMMA-acetone solution. The typical time to PMMA-pattern 1 scaffold array was 10 minutes. Similarly, double-sided tape was cut using a commercially available craft cutter **(Figure 2Aiii)**. We could robustly control the size of the well area by altering the pattern of PMMA printing to use larger diameter circles **(Figure 2Bi)** in the case of the 96-SPOT plate or by adjusting the line width via the use of different printing nozzle sizes **(Figure 2Bii)** in the case of the 384-SPOT plate design. Representative images of the 96-well and 384-well patterns are shown in **Figure 2Ci** and **2Cii**, where the PMMA-acetone solution contained blue dye to aid in visualization. Note that for the 96-SPOT pattern, in addition to a layer drawing an array of 96 circles, we also included a traveling salesperson (TSP) walk pattern to infiltrate exposed inter-well spaces using the PMMA-acetone solution **(SI Figure 1)**. This was an essential design component to ensure any slight misalignment during device assembly did not result in leakage between wells.

**Figure 2:**
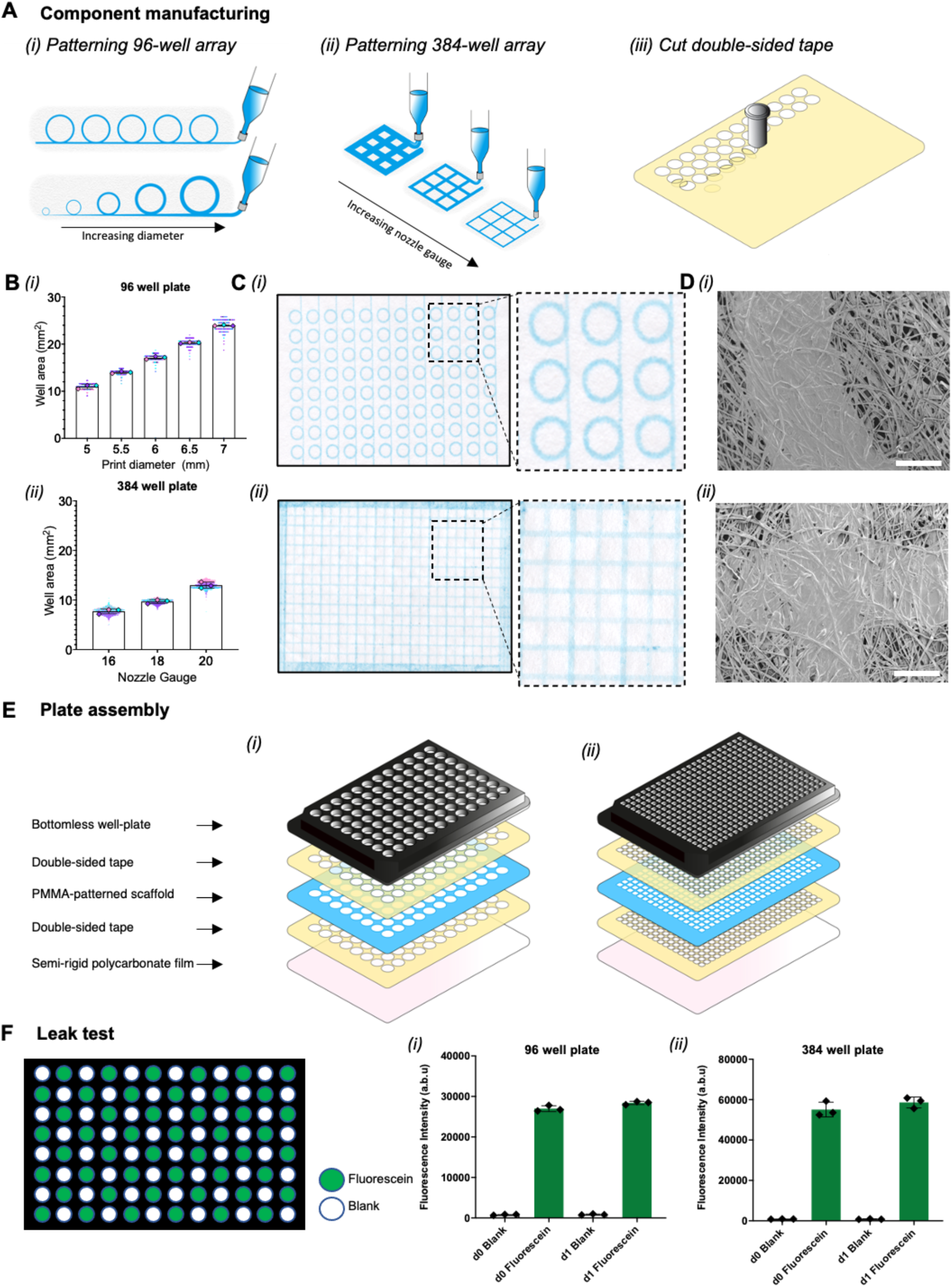
Polymer-infiltration facilitates the fabrication of a multi-well paper-based culture platform. **A)** Schematic representation of the manufacturing process for each SPOT component. (i) The PMMA-pattern for the 96-well scaffold array can be customized via the print pattern diameter and print speed, (ii) The PMMA-pattern for the 384-well scaffold array can be customized by altering the line width via the nozzle gauge. (iii) The double-sided tape is cut using a craft cutter to match the well-array. **B)** Quantification of the area of each well in the (i) 96-well plate indicated that well area correlated with the chosen print pattern diameter and the (ii) 384-well plate indicated that well area was defined by the line width (i.e. the selected nozzle gauge). Graphs show the mean ± SD of 3 independent experiments. **C)** Scanned representative image of a PMMA-infiltrated (blue) scaffold to create a (i) 96-well array and (ii) 384-well array. Image for 96-well array shown without TSP pattern in order to highlight the PMMA boundary generated with a single printed arc. **D**) Representative SEM image of a PMMA-infiltrated scaffold for a (i) 96-well plate and (ii) 384-well plate. Scale bars are 500µm. **E)** Schematic representation of the plate assembly for (i) 96-SPOT and (ii) 384-SPOT. **F)** 24-hour leak test experiment shows that fluorescein-containing wells (green) and blank PBS-containing wells (white) do not mix in the (i) 96-SPOT and (ii) 384-SPOT.

Following PMMA-patterning, the paper sheet was aligned to double-sided tape and adhered to a commercial no-bottom well plate and a thin optically transparent polycarbonate film. We selected an optically transparent material for the well bottom, to ensure the device would enable image-based readouts of cellular behaviour over time. To ensure that the PMMA barriers prevented any liquid leakage between wells, we confirmed that the PMMA polymer had infiltrated effectively into the paper-scaffold and blocked the scaffold pores **(Figure 2Di, 2Dii)** using scanning electron microscopy (SEM). After assembling the components of 96-SPOT **(Figure 2Ei)** and 384-SPOT **(Figure 2Eii)**, we also assessed liquid leakage between wells by performing a leak-test, adapted from methodology presented by the Whitesides’ and Lockett groups [44], [64] in which fluorescein is added to alternating wells of assembled 96-SPOT and 384-SPOT plates in a checkerboard pattern and fluorescence is measured on day 0 and day 1 **(Figure 2F)**. Over a 24 h period, we found that fluorescein did not travel into the blank wells for both the 96-SPOT **(Figure 2Fi)** and the 384-SPOT plate designs **(Figure 2Fii)**. We therefore were confident that our PMMA infiltration method provided a robust strategy to pattern the scaffold sheet into individual wells in which liquid transfer between wells did not occur.

### 3.2 Optimization of microgel formation in SPOT

After optimization of the fabrication of SPOT, we next set out to develop a reproducible method to load each scaffold well with cell-gel suspension to create a microtissue. We manually seeded 96/384-SPOT with GFP-expressing KP4 cells suspended in a collagen gel using a standard single-channel pipette. We then acquired widefield fluorescence images using an automated microscope and assessed the fluorescence variation within and between wells to assess the reproducibility of our manual seeding process. Representative images of the 96-SPOT array and 384-SPOT array are shown in **Figure 3A and 3B**, respectively. Without optimization of the seeding process, variation within wells can occur due to premature gelation of the prepared cell-gel solution and/or uneven seeding of the wells due to drying. Further, cell-gel aggregates can form due to poor cell-gel spreading or the tissues can lack cells due to improper pipetting of the viscous cell-gel solution. Both these features are problematic as cell aggregates can result in gradients of cell viability within the microtissue, while constructs lacking cells can result in inaccurate results in cell-based cytotoxicity assays. Furthermore, hydrogels are sensitive to temperature and humidity levels during seeding and gelation stages. In order to maintain humidity and temperature control during seeding, we added cold PBS to the spaces between the wells of the 96-SPOT. In the case of the 384-SPOT, we used a membrane to cover the wells during seeding and incorporated a humidification chamber to aid the manual seeding process.

**Figure 3:**
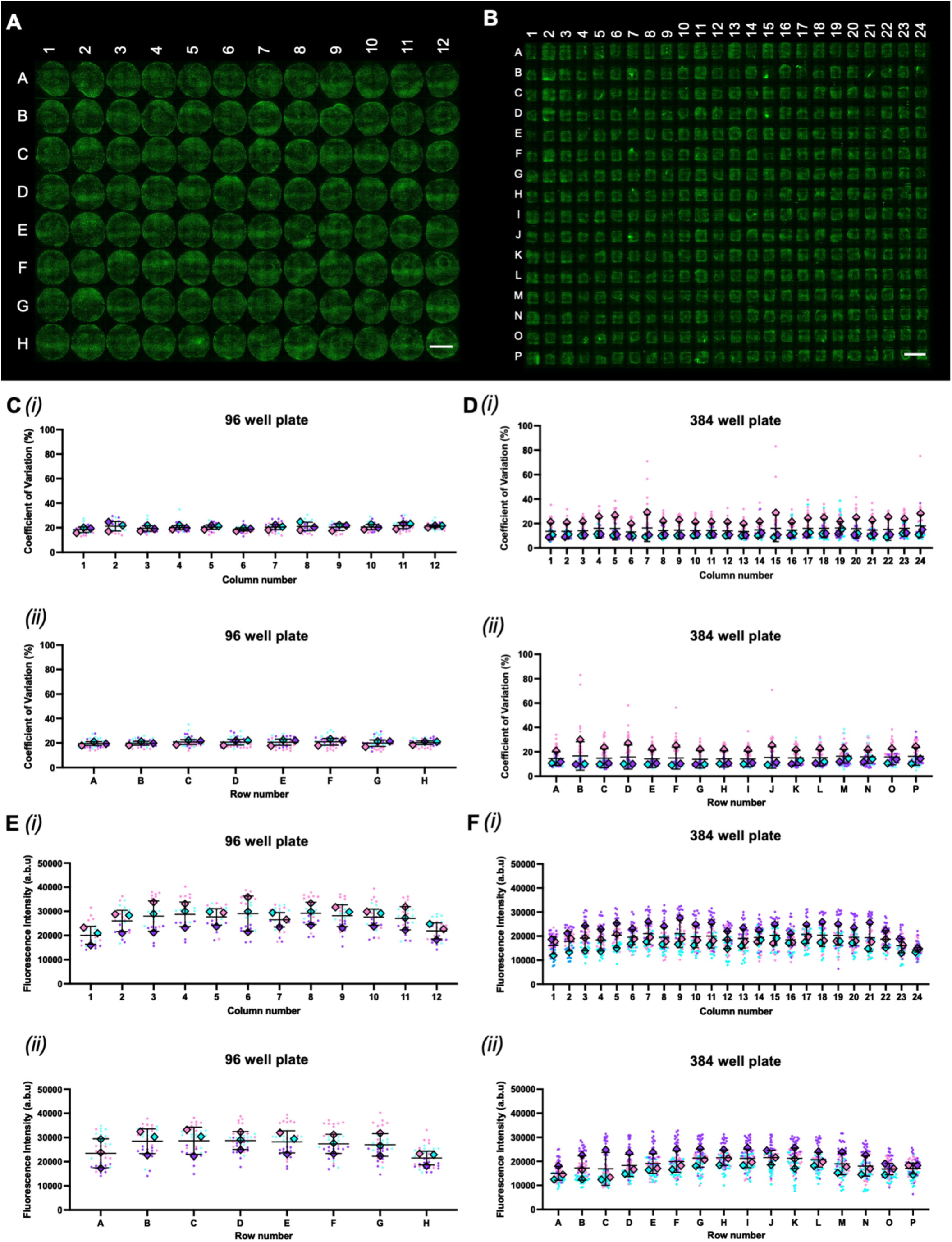
Assessment of fabrication variation for quality control of paper-supported tissue construct arrays. **A)** Representative widefield image showing 96-wells seeded with 30 × 10^6^ GFP-labelled (green) KP4 cells/mL collagen hydrogel. Scalebar is 4mm. **B)** Representative widefield image showing 384-wells seeded with 30 × 10^6^ GFP-labelled (green) KP4 cells/mL collagen hydrogel. Scalebar is 4mm. **C)** Assessment of coefficient of variation (SD/mean x 100%) measured from widefield images of a 96 well plate on day 0. Graph displays the mean coefficient of variation ± SD of 3 independent experiments. ANOVA revealed no significant differences between columns (i) or rows (ii). **D)** Assessment of coefficient of variation measured from widefield images of a 384 well plate on day 0. Mean ± SD of 3 independent experiments. ANOVA revealed no significant differences between columns (i) or rows (ii). **E)** The intrawell variation was measured by performing the alamarBlue® assay on 96-wells seeded with 30 × 10^6^ GFP-labelled KP4 cells/mL collagen hydrogel. Mean ± SD values are organized **i)** by column or **ii)** by row for 3 independent experiments. ANOVA revealed no significant differences between columns (i) or rows (ii). **F)** The intrawell variation in cell viability was measured by performing the alamarBlue® assay on 384-wells seeded with 30 × 10^6^ GFP-labelled KP4 cells/mL collagen hydrogel. Mean ± SD values are organized **i)** by column or **ii)** by row for 3 independent experiments. ANOVA revealed no significant differences between columns (i) or rows (ii).

To quantify the within (intra) well variation associated with our seeding process, we used an image-based script previously developed in [39]. Specifically, we first quantified the mean grey value (MGV) of fluorescence associated with each well containing the GFP-expressing KP4 cells. The use of MGV enabled identification of empty wells (i.e. low MGV) or wells containing excess gel (i.e. high MGV). Next, we measured the MGV of 100 randomly selected regions in each well to obtain the standard deviation (SD). Using the MGV of the well and the SD of 100 regions, we calculated the coefficient of variation as a percentage of the mean (i.e. %CV = MGV/SD x 100%). A high %CV corresponded to heterogeneous distribution of GFP signal within a well **(SI Figure 2)**. We used this metric to assess both the 96-SPOT plate **(Figure 3C)** and 384-SPOT plate **(Figure 3D)** and found no significant difference in the within well variation for wells in different (i) columns or (ii) rows. We also benchmarked the variation associated with a 384-SPOT plate against standard hydrogel plugs of different gel volumes in a 384-well plate **(SI Figure 3)**. The variation (%CV) between different wells was significantly lower than the variation associated with hydrogel plugs **(SI Figure 3A)**, likely due to the flat geometry of the scaffold suspended hydrogel in SPOT **(SI Figure 3B)** compared to the curved meniscus associated with a hydrogel plug **(SI Figure 3C)**.

We next compared the between (inter) well variation for 96-SPOT **(Figure 3E)** and 384-SPOT **(Figure 3F)** using an alamarBlue® assay. The alamarBlue® assay is a fluorometric method that detects changes in metabolic activity via mitochondrial reduction of resazurin to resorufin [68], [69]. While the alamarBlue® assay cannot provide spatially-resolved information within a well, we chose to use this standard assay to assess variation and confirm that our image-based metric was providing a reliable measurement of cell viability in SPOT. When comparing the alamarBlue® reduction between columns and between rows using one-way ANOVA, we found no statistically significant differences in either 96- or 384-SPOT. Based on this data, we concluded that we had established a robust SPOT seeding protocol with minimal within well and well-to-well variation that would allow us to reproducibly manufacture arrays of microtissues amenable for downstream cytotoxicity high-throughput assays.

### 3.3 Assessing chemotherapeutic response of a PDAC cell line in 96-SPOT

We next set out to demonstrate the use of the SPOT platform for response-to-therapy studies. To do this, we performed a proof-of-concept experiment to determine a drug dose-response relationship for gemcitabine monotherapy using the PDAC cell line KP4, following the experimental timeline shown in **Figure 4A**. First, we fabricated a 96-SPOT array containing GFP-labelled KP4 cells at a low cell density (i.e. 3 ×10^6^ cells/mL collagen hydrogel). The low cell density was chosen to accommodate the rapid growth of the KP4 cells and prevent overgrowth in the vehicle control conditions. After 24 h of culture in the SPOT array, we treated cells with a half-log serial dilution of gemcitabine, ranging from 0.001 - 100 µM. We then incubated the plate for 3 days before assessing cell viability using both images of the GFP-labelled cells and the alamarBlue® assay. Representative widefield 4x images of treated cells on day 3 are shown in **Figure 4B**. We generated dose-response curves from both imaging data **(Figure 4C)** and alamarBlue® measurements **(Figure 4D)**. While we observed slight differences between the dose response curves generated using each method, we found no significant differences between the IC_50_ observed from either assay over 3 independent experiments **(Figure 4E)**. This data indicates that our microscopy-based method to quantify response to therapy enabled accurate measurements of the IC_50_ value for gemcitabine-treated KP4 PDAC cells, that matched those made using a standard microplate-based readout using the alamarBlue® assay.

**Figure 4:**
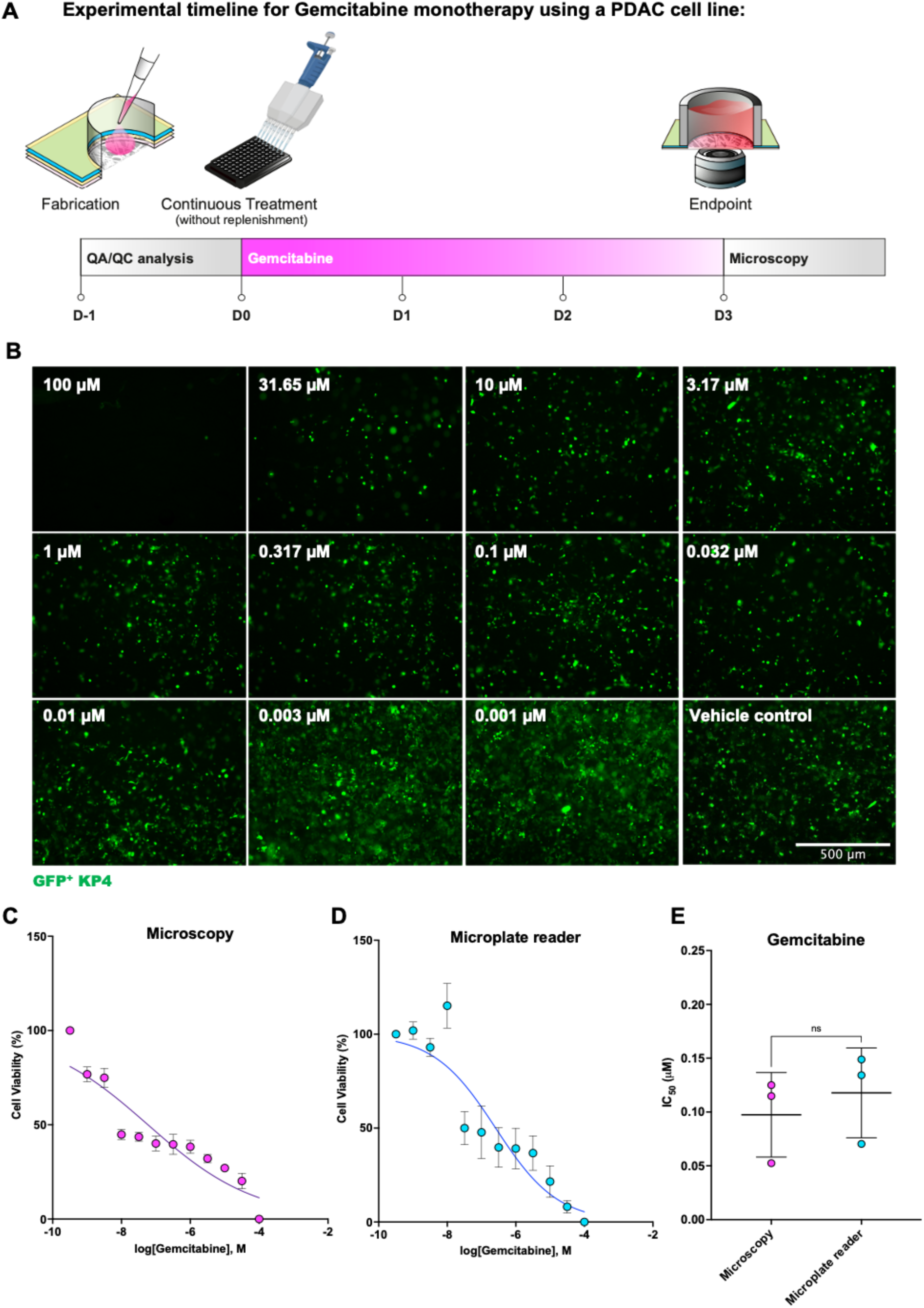
Assessing gemcitabine response of a PDAC cell line using 96-SPOT. **A)** Experimental timeline for gemcitabine monotherapy treatment experiments of the KP4 cell line. **B)** Representative widefield fluorescent images of GFP-labelled KP4 cells (green) associated with each gemcitabine concentration on day 3 at the end of drug treatment. Scalebar is 500µm. **C)** Half-log dose response curve for gemcitabine calculated using microscopy data. KP4 cell viability was measured using an ImageJ script which measures the mean gray value of the fluorescent image. Graph plots the mean viability ± SEM from 3 independent experiments. **D)** Half-log dose response curve for gemcitabine calculated using alamarBlue®. KP4 cell viability was measured using a standard alamarBlue® assay and microplate measurements. Graph plots the mean viability ± SEM of 3 independent experiments. **E)** Comparison of Gemcitabine IC_50_ measured using microscopy and microplate reader. IC_50_ values from both methods were not statistically different (t-test). Plot shows the mean IC_50_ ± SD from 3 independent experiments, i.e., IC_50, KP4, microscopy_ : 0.10 µM ± 0.04, IC_50, KP4, microplate reader_ : 0.12 µM ± 0.04.

### 3.4 Incorporation of patient-derived organoids in SPOT

Patient-derived organoids (PDOs) provide a more representative culture model of tumour heterogeneity and *in vivo* drug response, but are more challenging to incorporate into standard assays and/or platforms [32]. This is partially because they require 3D culture methods that are typically performed in hydrogel domes or plugs which are challenging to mass fabricate and/or image. Due to the suitability of SPOT to address these technical challenges, we set out to demonstrate the compatibility of PDOs in our 96-SPOT platform. We first assessed the growth of PDOs over 6-days in SPOT culture using widefield microscopy images **(Figure 5A)**. To do this we seeded 96-SPOT wells with a low density (3 ×10^6^ cells/mL collagen-Matrigel™ hydrogel) of GFP-expressing PDAC PDOs (specifically model PPTO.46) that had been dissociated to single cells and suspended in a 75% collagen-25% Matrigel™ hydrogel blend which has been reported previously by our group [39] for effective organoid culture. The images of the PDO-derived cells did not show elongated structures, suggesting that the paper fibers did not induce any significant morphological changes, which was consistent with previously reported observations [39].We then used MGV measurements of widefield fluorescence images to monitor growth. We found that the PDO-derived cells remained viable and grew over 6 days in 96-SPOT culture **(Figure 5B)**. This suggested that the chosen cell density (i.e. 3 ×10^6^ cells/mL collagen-Matrigel™ hydrogel) allowed for unrestricted growth over the 6-day experimental timeline.

**Figure 5:**
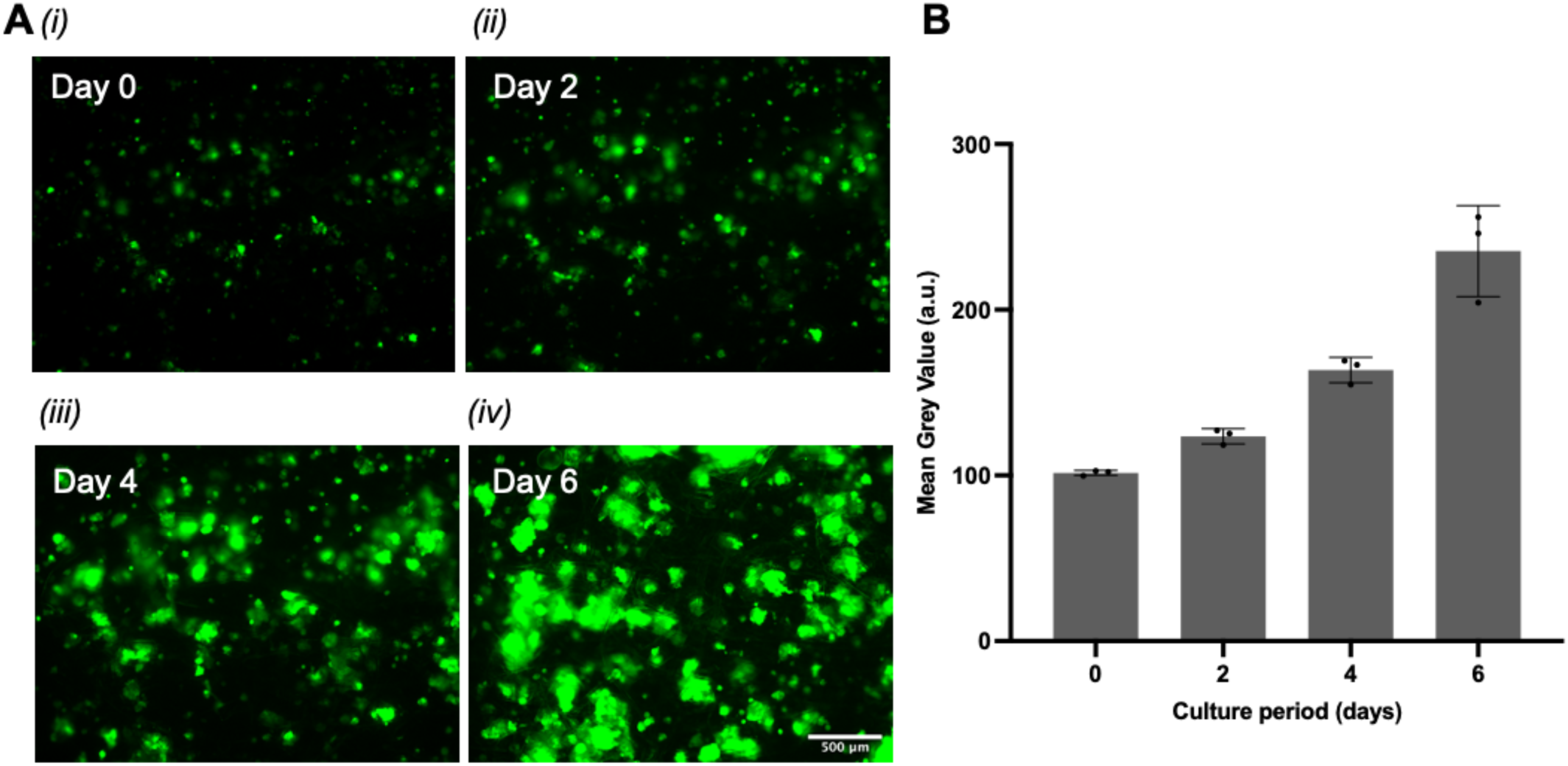
Organoid-derived PPTO.46 cells grow over time in 96-SPOT. **A)** Representative fluorescent image of GFP-labelled PPTO.46 cells (green) on (i) day 0, (i)iday 2, (iii)day 4, (iv)day 6. Scale bar is 500µm. **B)** Mean gray value calculated from PPTO.46 images. Graph plots the mean ± SD of 3 independent experiments.

### 3.5 Assessing chemotherapeutic response of PDAC PDOs in 96-SPOT

Having demonstrated the ability of PDOs to grow in SPOT, we next performed a dose-response assay in response to a 5-day treatment with gemcitabine following the experimental timeline shown in **Figure 6A**. On day -1, we seeded 96-SPOT with GFP-expressing PPTO.46 PDOs dissociated to single cells at a density of 3 ×10^6^ cells/mL of collagen-Matrigel™ hydrogel. After 24 h, we treated SPOT organoid cultures using a half-log serial dilution of gemcitabine, ranging from 0.0003-31.65 µM. Cultures were treated for 5 days and then imaged using an automated widefield microscope (**Figure 6B**). At this day 5 timepoint, cell viability was assessed using both microscopy images **(Figure 6C)** and the alamarBlue® assay **(Figure 6D)**. Similar to the previous experiment with the KP4 cell line, we observed slight differences between the microscopy-based and the microplate reader-based results. As observed in our experiments with cell lines however, we found no significant differences between the IC_50_ measured using either assay over 3 independent experiments **(Figure 6E**). These results highlight the accuracy of a microscopy-based readout for assessing PDO response to therapy in our SPOT platform.

**Figure 6:**
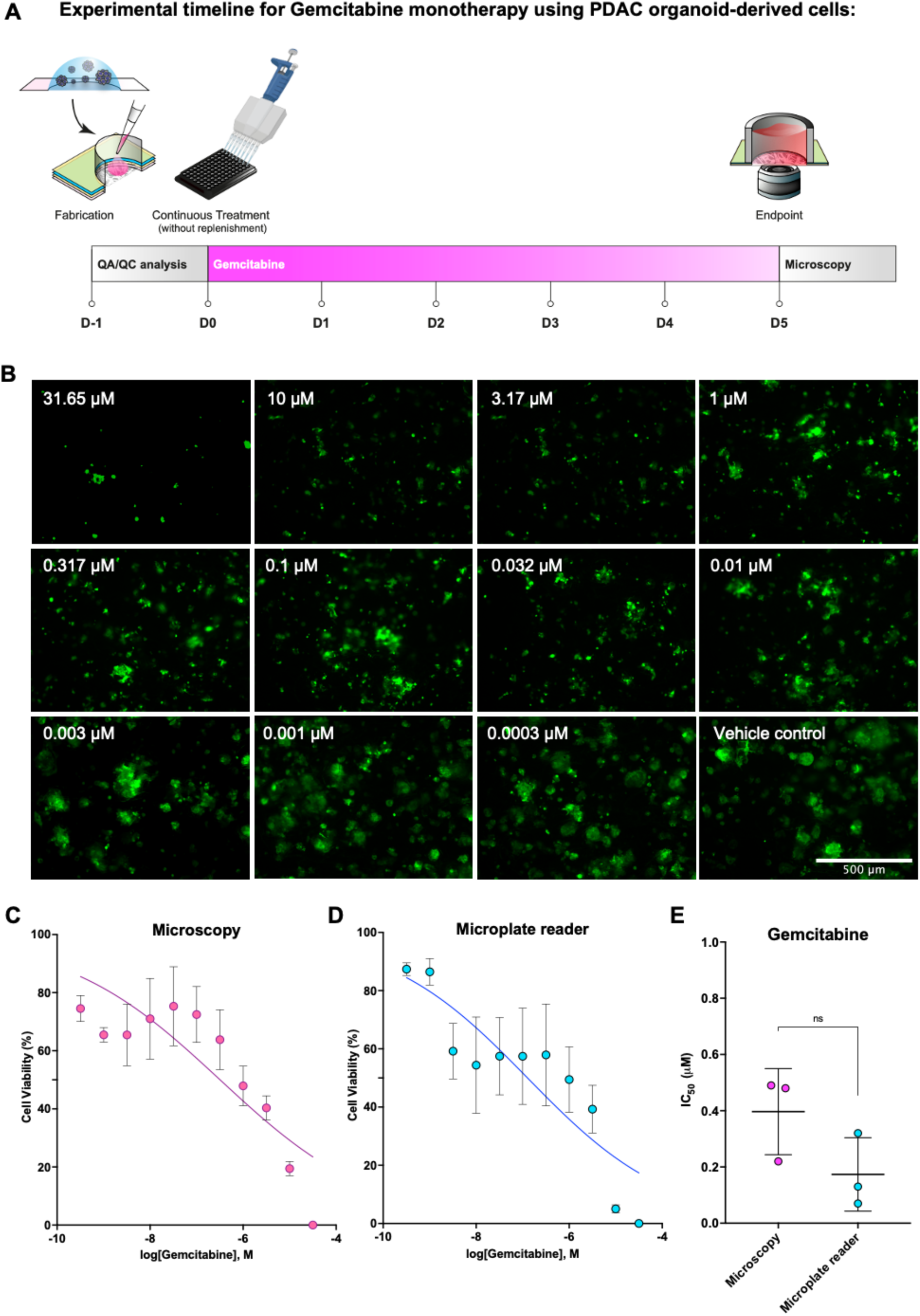
Assessing gemcitabine response of PDOs using 96-SPOT. **A)** Experimental timeline for gemcitabine monotherapy treatment experiments of PPTO.46 organoid-derived cells **B)** Representative widefield fluorescent images of GFP-labelled PPTO.46 cells (green) associated with each gemcitabine concentration after 5 days of treatment. Scalebar is 500µm. **C)** Half-log dose response curve for gemcitabine measured using microscopy images. PDO viability was measured using an ImageJ script which measures the mean gray value of the fluorescent image. Plot shows the mean viability ± SEM from 3 independent experiments. **D)** Half-log dose response curve for gemcitabine measured using a standard alamarBlue® assay. PDO viability was measured using an alamarBlue® assay and microplate measurements. Graph plots mean viability ± SEM from 3 independent experiments. **E)** Comparison of Gemcitabine IC_50_ measured using microscopy and microplate reader. IC_50_ values from both methods were not statistically different (t-test). Plot shows the mean IC_50_ ± SD from 3 independent experiments, i.e., IC_50, PDO, microscopy_ : 0.39 µM ± 0.15, IC_50, PDO, microplate reader_ : 0.17 µM ± 0.13.

### 3.6 Using 384-SPOT to scale-up PDAC PDO chemotherapeutic testing

Having demonstrated that we could incorporate PDAC PDOs into 96-SPOT and determine the gemcitabine drug-response curve, we next set out to demonstrate we could scale this assay using PDO cultures into 384-SPOT. Our drug response assay followed the same timeline as the one used in 96-SPOT and is shown in **Figure 7A**. On day -1, we seeded GFP-expressing PDOs dissociated to single cells at a cell density of 3 ×10^6^ cells/mL of collagen-Matrigel™ hydrogel into 384-SPOT. On day 0, we generated half-log serial dilutions of gemcitabine hydrochloride, 5-fluorouracil, paclitaxel, irinotecan hydrochloride (SN-38), and oxaliplatin. We then treated cells for 5 days before assessing the cell viability using both microscopy **(Figure 7B)** and the alamarBlue® assay **(Figure 7C)**. We found no significant differences between the IC_50_ observed from either assay over 3 independent experiments **(Figure 7D)**. Our data demonstrated that we were able to obtain IC_50_ values for each monotherapy, using half of a 384-SPOT plate for each replicate, highlighting the utility of 384-SPOT and image-based analysis to determine drug response of PDOs to several PDAC drugs in parallel.

**Figure 7:**
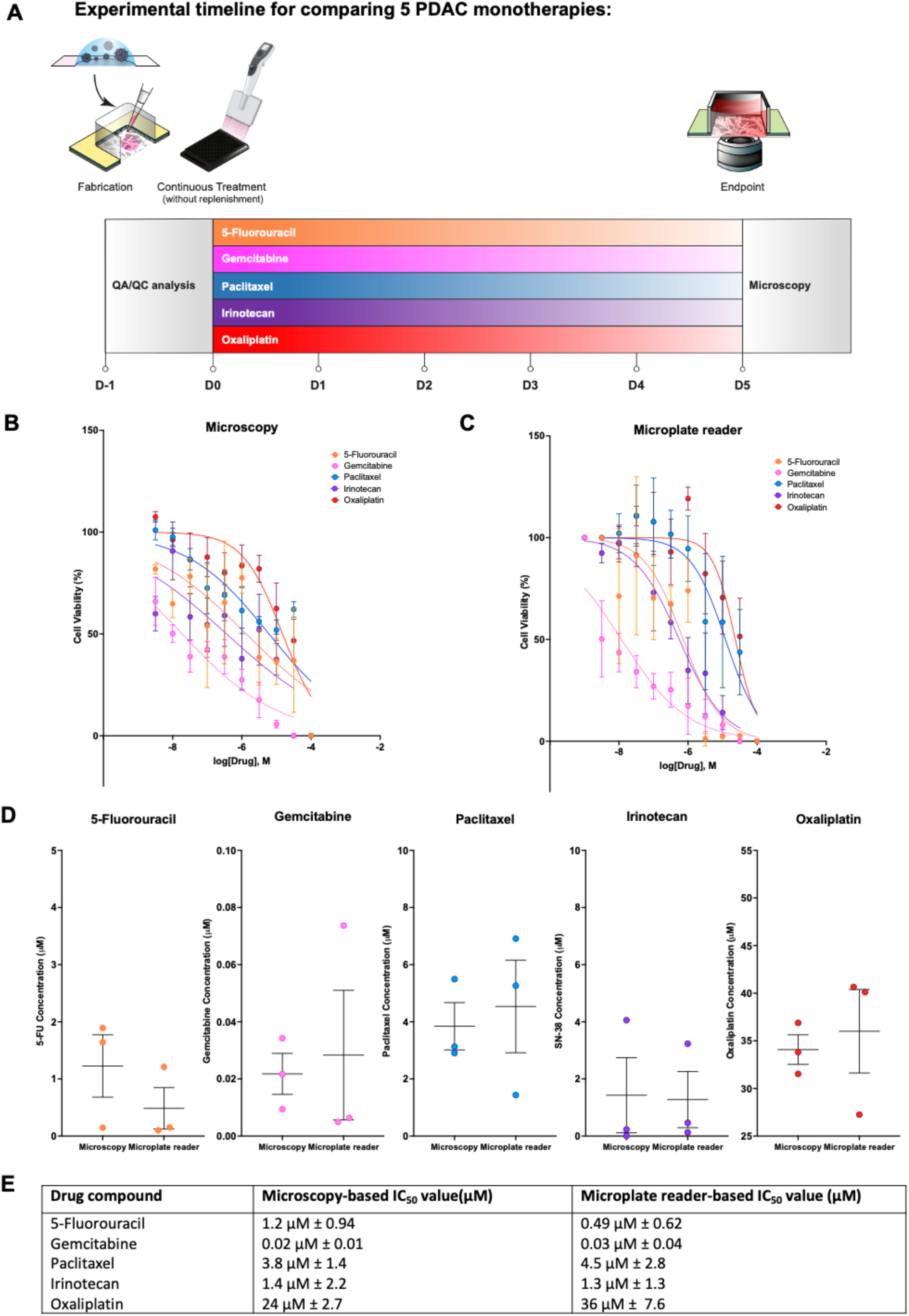
Assessing chemotherapeutic response of PDOs using 384-SPOT. **A)** Schematic of experimental timeline for 5 monotherapies of PPTO.46 organoid-derived cells **B)** Half-log dose response curve for 5 chemotherapeutic agents measured using microscopy images. PDO viability was measured using an ImageJ script which measures the mean gray value of the fluorescent image. Plots show the mean viability ± SEM from 3 independent experiments. **C)** Half-log dose response curve for 5 chemotherapeutic agents measured using standard alamarBlue® assay. PDO viability was measured using an alamarBlue® assay and microplate measurements. Graph plots the mean viability ± SEM from 3 independent experiments. **D)** Comparison of IC_50_ values measured using microscopy and microplate reader. IC_50_ values from both methods were not statistically different for each of the 5 chemotherapeutic agents. Plot shows the mean IC_50_ ± SEM from 3 independent experiments. **E)** Summary table showing IC_50_ values ± SD measured using microscopy and microplate reader.

## 4.0 Discussion

Drug discovery programs are beginning to exploit recent advances in patient-derived organoid cultures which have been shown to better model the heterogeneity observed within and between patients [23]. Approaches to screen the impact of various drugs on different organoid models to explore patient heterogeneity have primarily cultured organoids in 2D monolayers [29], [30], on the surface of hydrogel layers with dilute hydrogel in the surrounding media (the “slurry” method) [17], or within 3D hydrogel plugs [31]. Only the last of these approaches enables true 3D culture, in which cells are embedded within ECM. Hydrogel plugs however, are not ideal as a large volume of the hydrogel within the plug is associated with the meniscus that forms when the gel contacts the well edges. This can limit the number of compounds that can be tested if small amounts of sample material are available [25], [70]. Further, hydrogel plugs do not facilitate longitudinal *in situ* imaging of organoid phenotype due to the curved surface and high gel thickness associated with the hydrogel meniscus. The ability to perform longitudinal imaging of tumour cell cultures is important as we [35], [36] and others [25], [71], have shown the benefits of using this approach to provide a deeper understanding of organoid response to therapy. Here, we described the use of a thin paper scaffold to create supported hydrogel microtissues without a meniscus that enable the culture and long-term imaging of tumour organoids for analysis of drug response.

While multi-well paper-supported *in vitro* platforms have been previously developed to generate supported hydrogels to probe the cellular response to chemotherapeutics, the majority have focused on the use of bulk colorimetric or fluorometric assays which can be measured using a fluorescent scanner at the endpoint of the assay [44], [45], [64] as opposed to exploiting the geometrical templating of the paper scaffold to enable generation of a meniscus-free culture that enable image-based readouts of cell response to therapy. This is in part because previously reported paper-based systems typically require disassembly of the device for image analysis which dramatically limits scalability of the approach if using image-based readouts. We therefore designed the SPOT platform with the primary design criteria being not to require device disassembly for analysis, but rather to enable non-destructive *in situ* image-based readouts to quantify the drug-response and cellular phenotypes. Further, unlike previous iterations of paper-supported systems which require the assembly of the device after cell-gel seeding, we also prioritized the design criteria that the SPOT platform be prefabricated and stored for “off-the-shelf” use, to better facilitate adoption of the tool by the organoid community and future integration with automated pipetting systems to enable platform scaling for molecular screening. Also, to further facilitate tool adoption by the community, we designed the SPOT fabrication to use an open-source and inexpensive XY-plotter to pattern SPOT scaffold to enable users to easily customize the layout of the cell-containing regions of a plate to their specific needs.

Another advantage of SPOT is that in addition to defining the dimensions of the cell-gel structure, the scaffold also mechanically reinforces the hydrogel, allowing for both improved handling of mechanically fragile hydrogels, and the use of different and higher cell densities which more closely resemble *in vivo* tissue parenchyma. The majority of studies using PDOs cultured the cells in naturally-derived hydrogels such as Matrigel™ or collagen I, which are fragile and can often be aspirated or destroyed during liquid changes. The presence of scaffold provides mechanical reinforcement to hydrogel cultures, which could be particularly advantage for studies requiring frequent washing steps or culture media changes. Further, the high structural integrity combined with the flat geometry of the microtissue surface enables easy addition of additional cell types on demand, such as immune cells, on to the microtissues. Scaffold-supported microtissues also facilitates higher cell density cultures. In hydrogel-only cultures, typically, cell densities must be low (∼500 organoid-derived cells per 384-well) in order to avoid tissue contraction and distortion of the culture geometry during hydrogel polymerization or over the course of the culture period. Here, we generated SPOT tissues containing ∼4500 organoid-derived cells per 384-well with no observable changes in hydrogel dimensions. The scaffold-supported microtissues will also be useful for incorporating other cell types that are contractile and require stiff anchorage for their long-term culture. In a standard gel-only system such as the hydrogel plug, the absence of a reinforcing substrate can cause microtissues to detach from the culture well, thereby limiting the ability to perform long-term *in situ* imaging of the cells using standard widefield microscopy. For example, stromal cells such as cancer-associated-fibroblasts can cause tissue contraction if seeded at moderate cell densities in soft hydrogels such as Matrigel™ and collagen hydrogels. In these situations, the porous cellulose scaffold limits gel contraction [41] and maintains the structure of the microtissue. We note this could be a disadvantage for studying certain kinds of biology as the scaffold reinforcement will alter the local mechanical forces experienced by the contractile cells and may result in changes in morphology or cellular state. For this reason, we suspect SPOT is likely not appropriate for studies probing mechanobiology, and cellulose scaffold-free systems such as our GLAnCE platform [35], [36] may be better for such types of problems. Further, the presence of the scaffold may also be unfavourable for studies utilizing readouts that require bright-field optical imaging, due to the opacity of cellulose scaffolds.

We anticipate further development of 96/384-SPOT has the potential for use in personalized medicine. The low cellularity of primary patient samples, found in some tumour types such as PDAC, can be an obstacle for traditional drug screening approaches to decipher differences in chemotherapy response between patients. SPOT requires low volumes of cell-gel material (i.e., 5μL of 96-SPOT and 1.5μL for 384-SPOT) for experimental set-up and analysis, providing the opportunity to create microtissues from rare or specialized cell types extracted from patient biopsies. For example, previous work has demonstrated that PDOs originating from different PDAC patients can show preferential response to a particular combinatory chemotherapy [70]. SPOT could be useful for comparing multiple PDO lines and/or multiple therapeutic approaches, and by using non-destructive readouts, SPOT can enable higher-content or multiplexed experimental analysis for each patient. Towards this vision, future work will incorporate the use of automated pipetting systems to minimize variation, randomize treatment wells, and increase the overall throughput of the experimental pipeline.

## 5.0 Conclusions

96/384-SPOT provides a robust methodology to generate thin, dense, and flat PDO-containing microtissues for high-throughput cytotoxicity assays. Here, as an example, we demonstrated the use of SPOT to identify the drug sensitivity of PDOs and highlighted the ability of SPOT to enable image-based readouts of cell response to therapy. Specifically, we developed an image-based IC_50_ metric to quantify the presence of viable tumour cells after drug treatment and validated this image-based approach to measurements of IC_50_ made using the commonly used alamarBlue® cell viability assay. We anticipate that the ability to perform image-based readouts of cell function in 3D in an “off-the-shelf” high-throughput platform will enable 96/384-SPOT to become a valuable preclinical disease model for assessing drug sensitivity and other cell responses (for example, a change in morphology) for comparing or identifying novel therapeutics. Future work will focus on increasing the data content that can be extracted from images of SPOT and the automation of the SPOT manufacturing. The use of automated pipetting systems can minimize variation, allow the randomization of treatment wells, and increase the throughput of the overall experimental pipeline. Using an automated, user-independent workflow, we also envision the future application of the SPOT platform for personalized medicine approaches.

## Acknowledgements

The authors acknowledge technical assistance from Justin Pham, and Erik Jacques. Icons of multichannel pipettes in Figures 4, 6 and,7 were created with BioRender.com.

## Data and materials availability

The datasets generated during and analyzed in this study are available from the corresponding author upon reasonable request.

## Author contributions

Conceptualization: NTL, APM, NCW

Formal analysis: NTL, XL, YZ, MM

Funding acquisition: APM

Investigation: NTL, RC

Methodology: NTL, NCW, RC, JLC

Project administration: APM, NTL

Resources: APM

Software: RC, JLC, SL, YZ, MM, NTL

Supervision: APM, NTL

Visualization: APM, NTL, RR

Writing – Original Draft: NTL, NCW, APM

Writing – review & editing: All authors

