## Supplementary Information for "An off-the-shelf multi-well scaffold-supported platform for tumour organoid-based tissues"

**<u>Supplemental Tables:</u>**

**SI Table 1: The paper-based tissue model enables higher media volume to tissue volume ratio compared to hydrogel gel plug model**

| *Platform* | *Sample gel volume (µL)* | *Media working volume (µL)* | *Media volume: tissue volume ratio (Dimensionless)* |
|---|---|---|---|
| *96-well 2D monolayer* | N/A | 200 | N/A |
| *384-well 2D monolayer* | N/A | 100 | N/A |
| *96-well 3D paper-based tissue* | 5.0 | 200 | 40:1 |
| *384-well 3D paper-based tissue* | 1.5 | 50 | 40:1 |
| *96-well 3D gel plug* | 20 | 200 | 10:1 |
| *384-well 3D gel plug* | 10 | 100 | 10:1 |

**Supplemental Figures:**

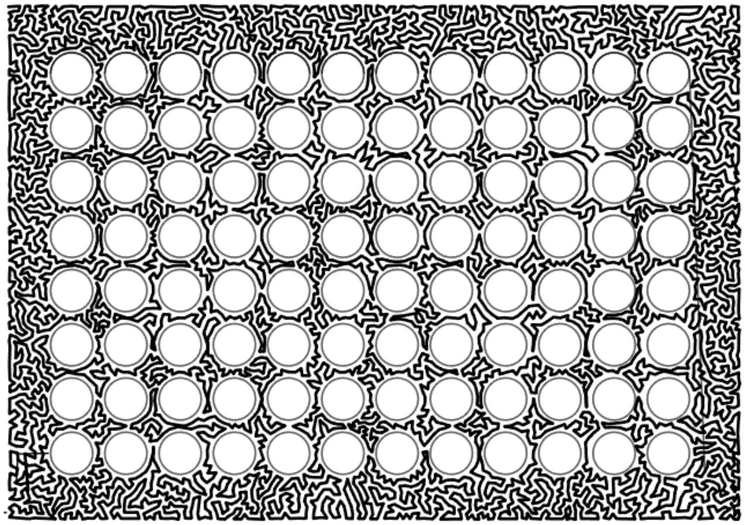

**SI Figure 1: The 96-well print pattern drawn using vector graphics software.**
Grey lines depict the pen-path which covers the surfaces between each well of a 96-well array. A custom traveling salesperson walk was created and imported into the vector graphics software, Inkscape, which is used to control the XY-plotter instrument.

A

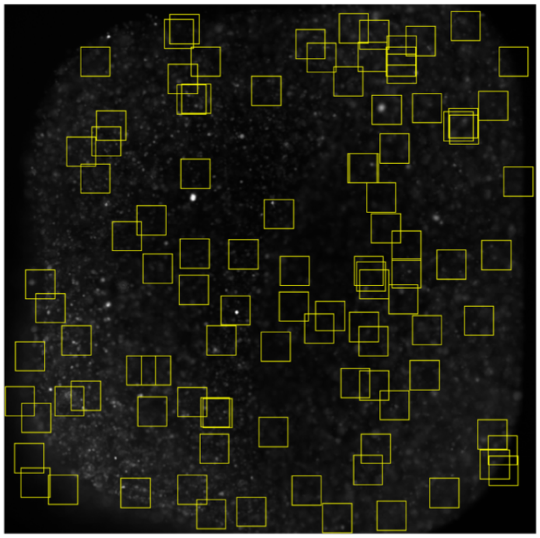

Sample mean grey value within 100 randomly-selected squares to obtain **standard deviation** within 1 well

B

| Image | Clump | No clump |
|---|---|---|
| Standard deviation | 83.6 | 57.1 |
| Mean | 218 | 278 |
| %CV | **38.4** | **20.5** |

C

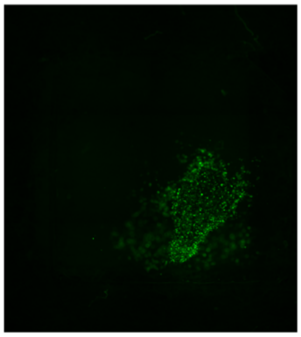

D

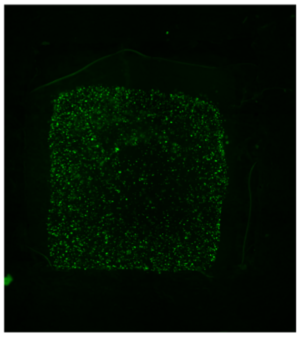

**SI Figure 2: Random sampling image algorithm for quality control assessment of seeded wells.** A) Representative image of 100 randomly distributed regions-of-interest (yellow square). B) Summary table showing image-based readouts for a well containing a C) clump or D) no clump. C) Representative full-well fluorescent image of a well containing a GFP-labelled KP4 cell aggregate (green). D) Representative full-well fluorescent image of a well containing GFP-labelled KP4 cells with no significant aggregates (green).

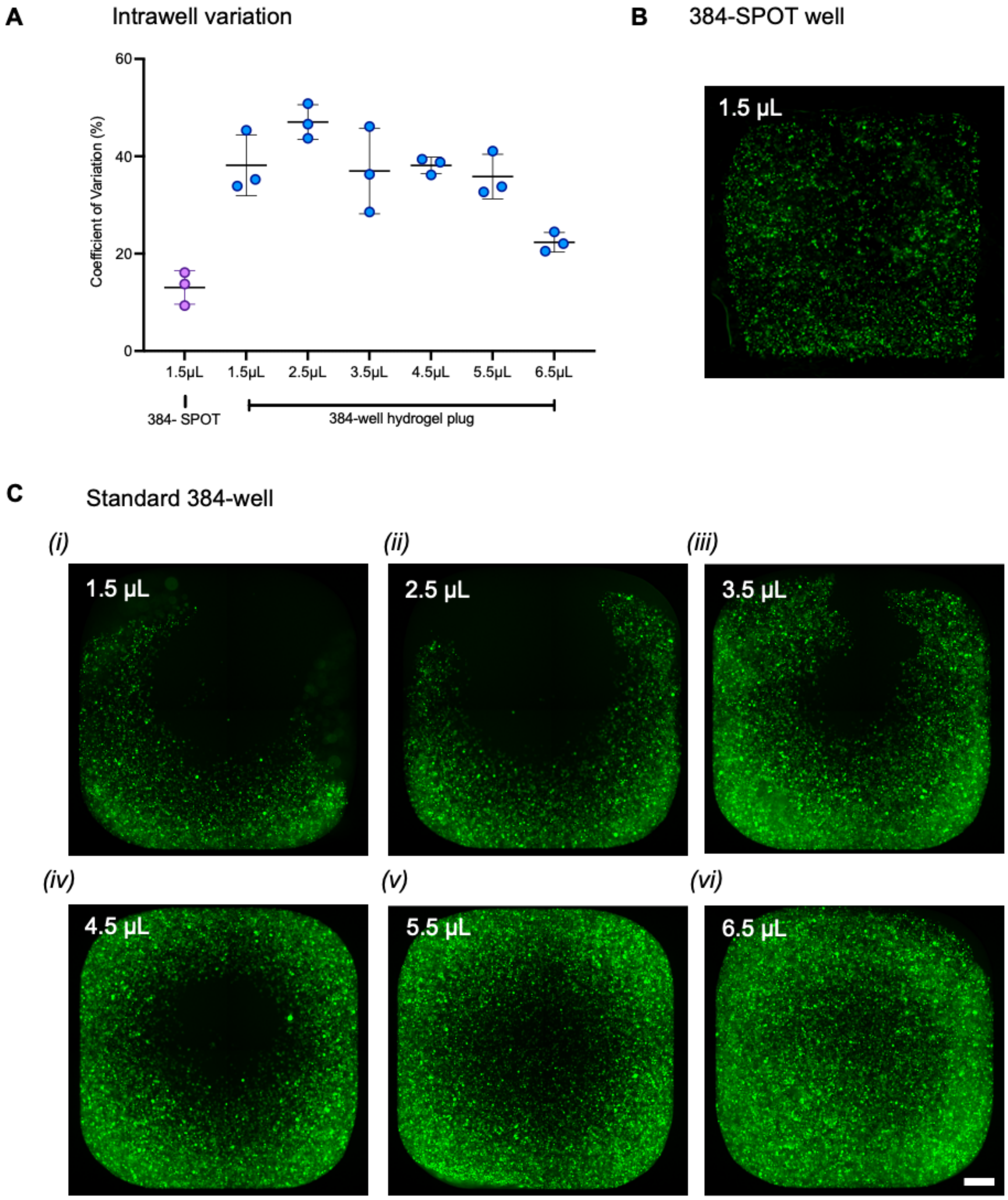


**SI Figure 3: 384-SPOT compared to standard 384-well hydrogel plugs**
**A)** Coefficient of variation percentage measured from acquired widefield images on day 0. Mean ± SEM of 3 technical replicates. **B)** Representative full-well image of one 384-SPOT well seeded with 1.5μL of collagen hydrogel containing 4.5 x $10^4$ GFP-labelled KP4 cells (green). **C)** Representative full-well image of a hydrogel plug in a standard 384-well seeded with **(i)** 1.5 μL, **(ii)** 2.5 μL, **(iii)** 3.5 μL, **(iv)** 4.5 μL, **(v)** 5.5 μL and **(vi)** 6.5 μL of collagen hydrogel containing 4.5 x $10^4$ GFP-labelled KP4 cells (green).
